# bgnorm: A Generative Statistical Framework for Background Correction, Normalisation, and Quality Control in Multiplex Spatial Proteomics

**DOI:** 10.64898/2026.08.05.743141

**Authors:** Malvika Kharbanda, Rafael Tubelleza, Yuqi Tan, Chin Wee Tan, Cooper Janke, Ismail Sebina, Gabrielle T Belz, Arutha Kulasinghe, Agus Salim, Dharmesh D. Bhuva

## Abstract

Multiplex spatial proteomics enables highly multiplexed in situ profiling but remains limited by technical variation arising from autofluorescence, non-specific antibody binding, instrument noise, and staining variability, affecting downstream biological tasks like cell typing. We present *bgnorm*, a statistical framework that describes fluorescence measurements using a generative mixture model of background, non-specific binding, and biological signal components. As natural statistical consequences, the model yielded three new methods: a background-correction method through probabilistic deconvolution of protein intensities, quality control metrics, and a quantile normalisation approach to unify measurements across markers, samples, and sequential slices. Across multiple multiplex imaging technologies, *bgnorm* improves signal separation and downstream marker positivity classification compared with existing preprocessing approaches. In expert-annotated datasets comprising over 406,000 marker positivity annotations, *bgnorm* achieved the highest classification performance and enabled accurate use of a single global positivity threshold across markers and samples. The method is implemented in the *bgnormR* and *bgnormpy* packages.

## Introduction

Spatial proteomics has emerged as a transformative approach for mapping protein localisation within intact tissue architecture, providing insight into cellular phenotypes, tissue organisation, and tumour microenvironment interactions that cannot be captured using bulk or dissociated assays^1, 2^. Over the past decade, spatial proteomics technologies have evolved from low-plex immunohistochemistry (IHC) to highly multiplexed imaging platforms including Akoya PhenoCycler-Fusion (PCF; formerly CODEX)^3^, Cell DIVE^4^ (Leica Microsystems, formerly GEHC), imaging mass cytometry (IMC)^5^, multiplexed ion beam imaging (MIBI)^6^, CyCIF^7^, and CosMx CellScape PSP^7^, among others, enabling simultaneous measurement of tens to hundreds of proteins across intact tissue sections. These technologies have advanced the study of tumour-immune microenvironment, tumour heterogeneity, and in situ cell-cell interactions across a wide range of diseases^8^. More recently, three-dimensional (3D) reconstruction approaches using serial tissue sections have extended these analyses beyond two-dimensional tissue architecture, enabling spatial profiling across entire tissue volumes^9^. Technological progress combined with the potential to uncover previously unexplored biology has resulted in the technology being crowned the method of the year in 2024^10^.

Despite these advances and the excitement in the field, quantitative analysis of multiplex immunofluorescence (mIF) imaging data remains a challenge due to poor characterisation of technical variation^11^. Investigations into the generative processes underpinning measured intensities remain unexplored resulting in a poor understanding into the sources of technical and biological variation. Raw fluorescence intensities represent a complex combination of true biological signal, tissue autofluorescence, non-specific antibody binding, detector noise, staining variability, and field-of-view effects. Unwanted sources of variation distort continuous intensity distributions, complicating downstream analyses including marker positivity classification, clustering, cell phenotyping, and spatial neighbourhood inference^12, 13^. While numerous approaches have been developed to process intensity data into biological insights, key steps of the pipeline, including normalisation, quality control, and marker positivity classification rely on heuristic approaches due to the lack of understanding into the measurement process of fluorescence imaging intensities.

Several normalisation approaches have been proposed to address technical variability in mIF data^14^. These methods broadly fall into two categories. The first are transformations that stabilise the variance, such as the log and inverse hyperbolic sine (sinh^−1^) transformations, which aim to reduce skewness in the distribution of protein intensities and to stabilise the variances making them less dependent on the mean. These methods correct the skewness in measurements but do not directly enable harmonisation of measurements across samples and proteins. They are often but not always coupled with the second category of methods that apply a global scaling to the intensity values of all cells with the aim of harmonising the marker intensity distributions through rescaling at the protein level, sample level, or both. These methods include approaches such as z-score normalisation^11^, ComBat^15^, centred log-ratio (CLR) transformation^16^, UniFORM^17^, percentile-based scaling^11^, mean normalisation, and min-max scaling. While methods based on descriptive summaries (e.g., means and percentiles) are generally non-parametric, z-score, ComBat, and CLR are parametric and impose invalid assumptions for immunofluorescence data, resulting in poor performance in recent benchmarks^17^. Intensity data are known to be multimodal^4, 18^ therefore the unimodal Gaussian assumption imposed by z-transformations and ComBat is violated. CLR is a transformation designed for compositional data, which intensity data is not since each marker is profiled independently, and the measurement of one protein is independent of other proteins in mIF assays. A common limitation of all global scaling methods is that all measurements are scaled equally, therefore erroneous measurements (e.g., intensity from non-specific antibody binding events) are scaled in the same way as positive signals.

Preprocessing impacts downstream analysis including cell typing which often relies on marker positivity-classification. Computational methods such as Otsu thresholding^19^, k-means clustering, and mixture model-based gating approaches^20^ are commonly used to assess marker positivity for intensities summarised at the cell level where the task is a binary classification of marker positivity. While deep learning frameworks attempt to address the normalisation problem through data augmentation strategies, studies have shown that normalisation of data prior to learning improves generalisation, specifically for cross-site datasets^21–24^. Indeed, a recent deep learning approach for marker positivity prediction includes data normalisation to harmonise datasets across sites, cohorts, and technologies ^25^. Together, these limitations highlight the need for a modelling framework for fluorescence imaging intensity data that explicitly accounts for the different sources of background-associated variation and that can be studied to develop statistical procedures to tackle key preprocessing tasks in fluorescence imaging data.

To this end, we present *bgnorm*, a modelling framework that identifies the different sources of intensity variation in fluorescence imaging data and uses a generative statistical model of the data to derive a background correction procedure, a quantile normalisation approach, and quality control metrics. While the original approach is designed for pixel-level data, we also propose a variant that normalises data summarised at the cell-level. We show that the cases where the model does not fit the data well enough are associated with quality issues, thus also functioning as a quality control approach whilst simultaneously affirming the modelling framework. We show that spatial variation in model parameters is minimal across whole slide images therefore the same approach can be applied for tissue microarrays and whole slide images without the need for spatial constraints. Finally, we demonstrate comparable performance of the pixel-and cell-level approaches, and the superior performance of *bgnorm* compared to commonly used methods through an extensive benchmark using an expert annotated multiplex imaging dataset. Since *bgnorm* enables intensity comparisons across samples, proteins, and 3D slices while independently normalising each image, it holds promise for the sequential normalisation of future atlas-scale spatial proteomic datasets. The proposed modelling framework can be used to develop future studies regarding additional properties of immunofluorescence data including differential intensity analysis.

## Results

### A mixture of log-normal convolutions to model and adjust fluorescence intensity measurements

The observation that fluorescence intensity data is a mixture of log-normal distributions has been made across various studies in the past and have resulted in the use of Gaussian Mixture Models (GMMs) to model log-intensities ^18, 26^. Their use has been limited to the study of cell-level log-intensities with the objective of identifying cells that are positive for a given marker. While the data display multiple peaks (Figure 1A), the number of mixtures and their interpretation is lacking, thus preventing downstream applications. As such, we began by studying the distribution of log-intensities following preprocessing to reduce the impact of very low intensity noisy signals (see Methods) in a Non-Small Cell Lung Cancer (NSCLC) dataset with 45 proteins measured across 42 patient samples.

**Figure 1:**
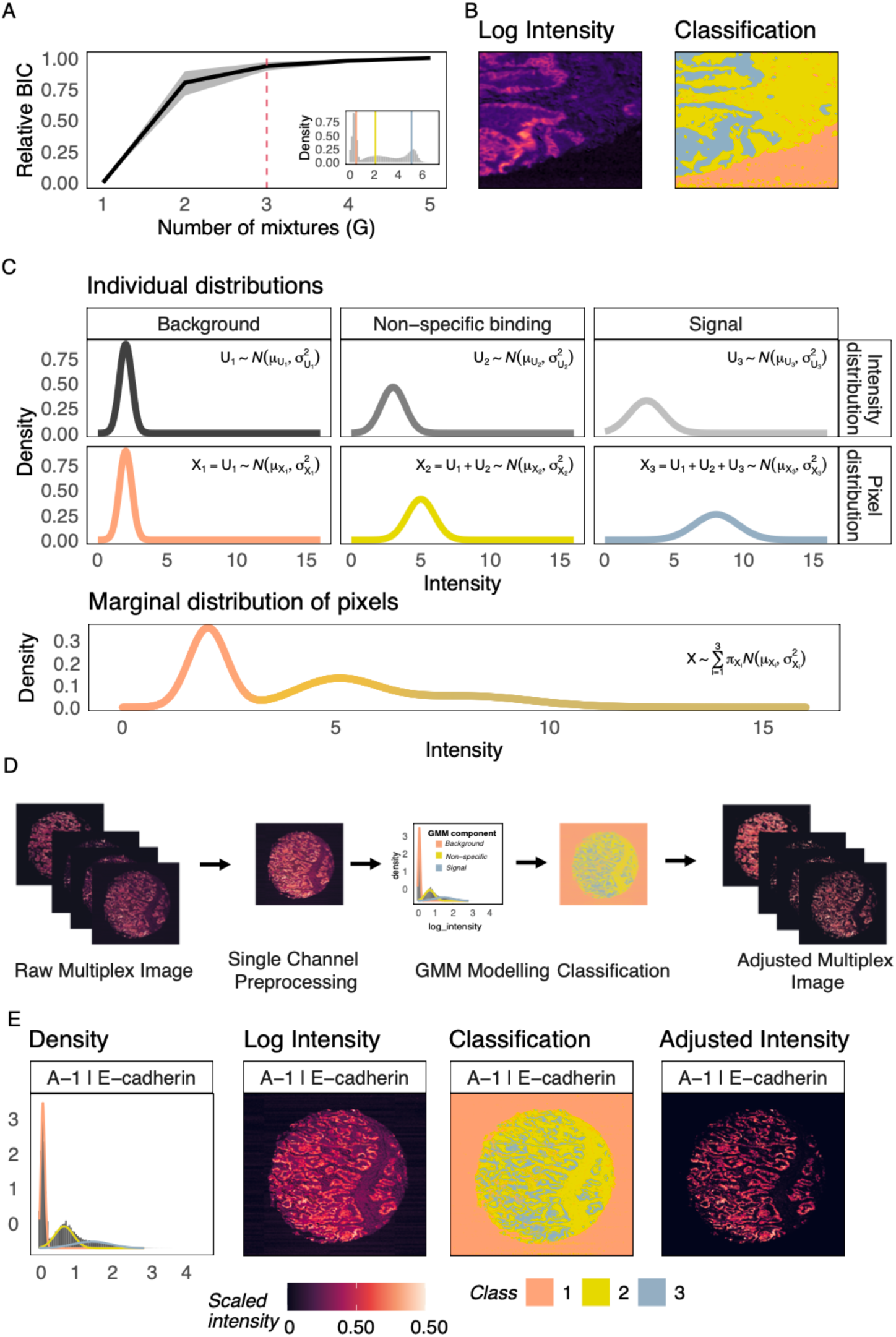
Model-based background normalisation of multiplex imaging data using bgnorm. A) Model-selection using the negative Bayesian Information Criterion (BIC) relative to the -BIC of a one-component model shows that a three-component mode (inset)l is appropriate for imaging data. B) Classification of pixels annotates the three components as background (orange), non-specific binding (yellow), and signal (blue). C) The bgnorm model assumes there are three dominant sources of illumination: background (U_1_), non-specific binding (U_2_), and biological signal (U_3_) modelled as lognormal distributions. Pixel intensities are modelled as successive convolutions X_1_, X_2_, and X_3_, resulting in the final marginal distribution which is a mixture of lognormal convolutions. (D) Overall schematic of the bgnorm algorithm with deconvolution of the signal component to obtain background-corrected intensities. (E) Application of bgnorm to an E-cadherin image from a non-small cell lung cancer (NSCLC) patient shows that three distinct components are identified from the data. Raw images show E-cadherin staining; however, contrast is low due to luminance in stromal regions. GMM accurately identifies three distinct spatial regions which allows bgnorm to adjust background effects and produce a high-contrast background adjusted image.

Gaussian mixture models with 1 to 5 components were fit to each protein across each core, and the Bayesian Information Criterion (BIC) was measured to determine the ideal number of components to represent the data. The relative BIC of each model compared to that of the one-component model showed that a three-component model was adequate for almost all proteins measured across all cores, with marginal gains offered by more complex models (Figure 1A). Studying the component allocation of pixels using the three-component model revealed that the three components coincided with distinct biological regions: i) background space with no tissue, ii) tissue regions lacking the protein of interest, and iii) tissue regions containing the protein measured (Figure 1B).

Combining this information with the different sources of intensity in fluorescence microscopy such as baseline instrument intensity, autofluorescence, non-specific binding, and target binding, we postulated a mixture of log-normal convolutions. Sources of intensity were ordered based on their expected intensity as: (i) low-intensity background signal, (ii) intermediate non-specific binding and autofluorescence, and (iii) high-intensity biological signal (top row of Figure 1C) and modelled as log-normal distributions with distinct means and variances. Each pixel is then a measurement of the sources of intensities it overlays. Pixels from regions with no tissue (blue in Figure 1B) only experience the background intensity, those from regions with tissue but without the protein of interest (yellow in Figure 1B) experience background intensity and intensities from non-specific binding events and autofluorescence, and finally pixels from regions with positive staining (red in Figure 1B) experience all sources of intensities, producing a successive convolution of the sources of intensity (middle row of Figure 1C). If the identity of the pixel was known, their distribution would be a convolution of log-normal distributions, however, since we do not have this information when studying the raw data, the marginal distribution of each pixel is a mixture of log-normal convolutions (bottom row of Figure 1C). This reasoning allowed us to develop a generative model of the fluorescence intensities, which was then used to deconvolve the signal component to compute background-corrected intensity measurements, thus leading to the *bgnorm* algorithm (Figure 1D).

Applying *bgnorm* to the E-cadherin image channel for one of the cores showed that the GMM fit for the data resulted in the identification of the three components, each representing background, non-specific binding and autofluorescence, and signal, with the latter identifying the epithelial cancerous tissue in this sample (Figure 1E). Model parameters were then used to compute background adjusted intensities, resulting in the suppression of intensities within the stromal regions (class 3 in Figure 1E), producing an image with sharper contrast for E-cadherin staining.

Raw intensity images are often unavailable, and the only accessible data would be cell-level intensity summaries (mean or median). To address this, we also propose an approximation of *bgnorm* that fits a two-component GMM representing autofluorescence/non-specific binding events and signal events. Data are still normalised using the deconvolution approach since the parameters required are available from the two components. We note that this approach is not a true reflection of a cell-level model since it assumes all or at least a majority of pixels in a cell belong to a single component which is seldom true, especially for membrane bound markers. Nonetheless, it offers a useful approximation for cases where data access is limited.

Background correction using *bgnorm* is performed for each sample and each marker independently, therefore comparisons across samples, markers and sequential slices of 3D datasets are not directly addressed. The modelling framework can be used to assess the degree of variability in these scenarios as we can study the model parameters directly. We evaluated variation in intensities across samples, markers, and 3D slices using a 3D multiplex skin imaging dataset containing 26 slices and 2 tissue regions profiled using the Cell DIVE platform^9^. A *bgnorm* model was fit to each marker in each slice across each sample and the model parameters were studied. Variation in the means for each component was marker specific with variation in the mean minimal across slices and samples for markers such as PCAD (Figure 2A). In contrast, significant slice-specific variation was observed for DDB2 for both the signal and non-specific binding/autofluorescence components (Figure 2A). Finally, markers such as AE1 exhibited a sample-specific difference in means with the global mean of the signal component across slices lower in one sample compared to the other (Figure 2A).

**Figure 2:**
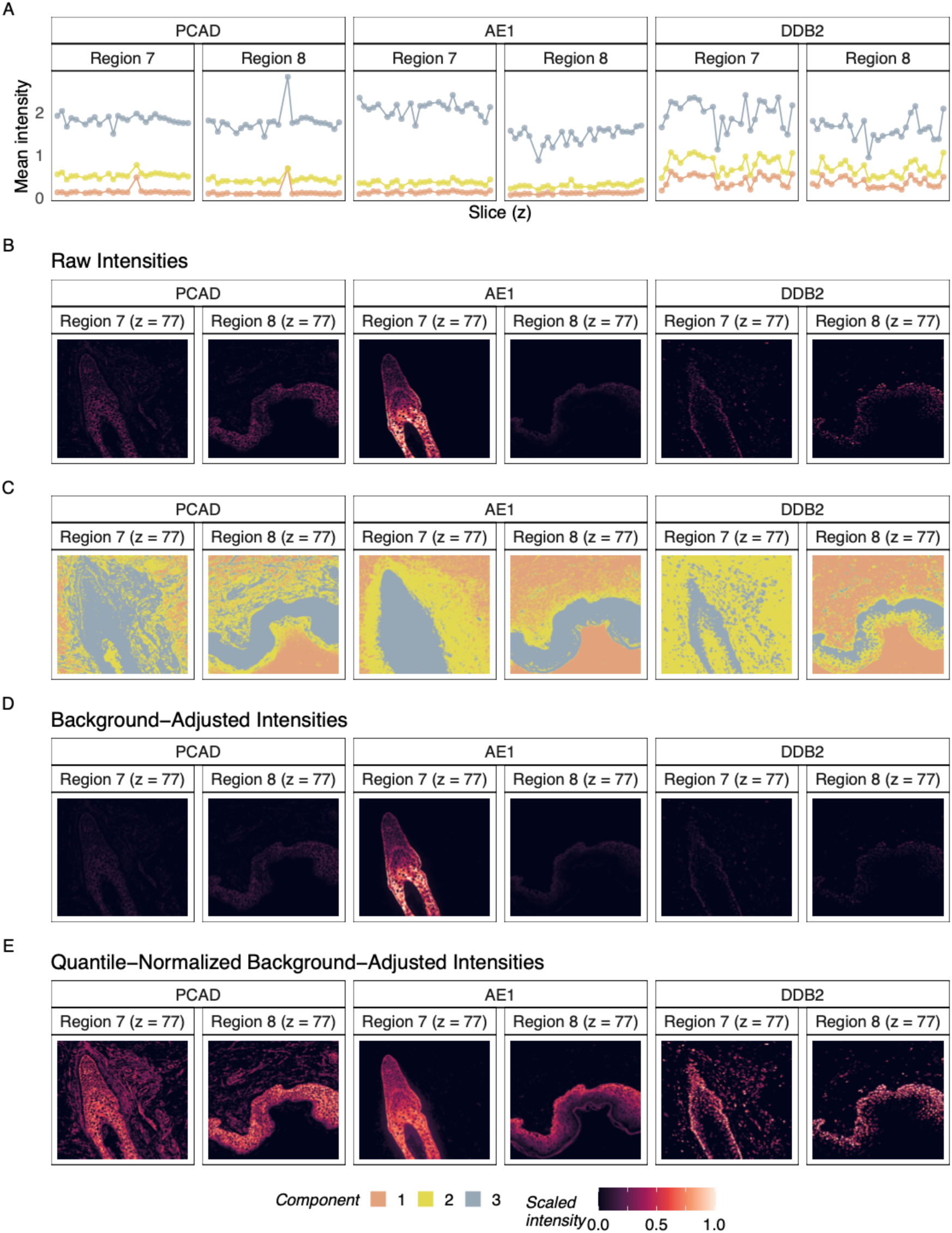
Quantile normalisation equalises the dynamic range of intensities across samples, slices, and channels. (A) Gaussian mixture model (GMM) component means across 3D slices for representative markers PCAD, AE1, and DDB2 for two tissue regions showing minimal variation, sample-level variation, and slice-level variation respectively. Spatial plots showing B) Raw log transformed intensities, C) Region classification, D) background-adjusted intensities, and E) quantile-normalised background-adjusted intensities, for a representative slice from 2 samples demonstrating cross-sample, cross-sample, and cross-channel comparability following quantile normalisation.

As expected, means of the background component were generally comparable across slices, samples, and markers as this variation stems from the instrument and the acquisition parameters. Variation in the means of the non-specific binding/autofluorescence component was marker-specific (Figure 2A), reflecting a difference in the binding affinities of antibodies. Differences in the means of the signal component for a given marker across samples or slices may reflect true biological differences in the abundance of the measured protein or technical differences in sample preparation that resulted in better staining in one sample compared to another. As it is difficult to differentiate between these two sources of variation, we proposed an independent quantile normalisation step that could be performed following background adjustment to each fluorescence image. In most cases, this additional step is recommended as it unifies measurements across samples, slices, and channels (Figure 2B), and should only be disabled if there is sufficient evidence of a biologically motivated difference in intensities (e.g., differences in metabolic markers across disease states). Unlike conventional sample-based quantiles, we computed a model-based quantile that used the background-adjusted quantile of the fitted signal component to rescale the data.

Studying a representative slice from both samples across the three markers PCAD, AE1, and DDB2, we saw that raw intensities were not comparable across images, particularly due to the higher intensity of AE1 in a single sample (Figure 2B). While *bgnorm* is able to identify the regions of signal (Figure 2C) and adjust for background effects, adjusted data are not comparable similar to the raw data (Figure 2D). Subsequent quantile normalisation brings the data to a unified scale, thus equalising intensities across markers, samples, and slices (Figure 2E). Since intensities are comparable across images, a single processing pipeline suffices in the analysis of all channels, slices, and samples.

Cell typing using spatial proteomic data usually involves assessing marker positivity for each cell and subsequently using knowledge of combinations of cell type markers to allocate identity to cells. A crucial step in this workflow is to assess marker positivity, which is subject to prior normalisation and the classification algorithm. Various methods have been developed for intensity normalisation, including *bgnorm*, while common choices for the binary clustering task are Otsu thresholding, GMM-based clustering, k-means clustering, and applying a fixed threshold to the data. Fixed thresholding is rarely performed as the data are on different scales across samples, slices, and markers, however, *bgnorm* rescales data to enable a uniform comparison thereby making fixed thresholding feasible. To evaluate the impact of *bgnorm* on downstream marker positivity classification, we benchmarked the method against commonly used normalisation approaches (see Methods) using three CODEX colon fields of view (FOVs) from the expert-annotated Pan-M multiplex imaging data composed of 406,352 gold-standard positivity labels ^25^. The combination of all normalisation methods and classification methods was tested to enable a fair comparison with F1 scores, precision, and recall measurements used to evaluate performance. Since performance depends on the quality of the data, which is protein-specific due to differences in antibody affinities, we assess performance at the protein-level. This is done by aggregating F1 scores into a median F1 score per method for each protein, and subsequently ranking method performance for each protein.

F1 scores showed that the pixel-level and cell-level variants of *bgnorm* with or without quantile normalisation outperformed all other normalisation methods across all datasets and markers (Figure 3A). In the context of determining marker-positive cells, the best *bgnorm* performance produced a mean F1 score of 0.64 which is 5.0% higher than the next best performing alternative method, *min-max* normalisation (mean F1 = 0.61). Interestingly, performance was comparable for the pixel-and cell-level variants of *bgnorm*, demonstrating that the cell-level approximation was effective. The three worst performing normalisation methods involved a standard *log10* transformation (mean F1 score of ∼0.33). This differs from the transformation *bgnorm* uses which borrows the idea of using a cofactor from the *arcsinh* transformation to suppress very low intensity noisy measurements, highlighting the importance of this preprocessing step. Classification using GMMs was generally worse than with Otsu thresholding or k-means clustering, likely due to the non-parametric nature of the latter two approaches. Coupling *bgnorm* with GMM-based clustering yielded the worst performance for *bgnorm*, however, this was expected since adjusted intensities are no longer a mixture of Gaussian distributions. In some cases, a marker was not expressed in most or all cells and this can be crucial in annotating cell types as it is the combination of presence and absence of markers that determines cell identity. As such, we assessed F1 scores in determining marker negativity as well (Figure3B) and this showed that most methods performed well in negativity classification with the worst performing being the log-transformation-based, CLR, and Arcsinh. Comparing performance across both classification tasks, we saw that *bgnorm* balanced performance across both tasks whereas methods like UniFORM and Double Z sacrificed positivity annotation for improved negativity classification (Figure 3C).

**Figure 3.**
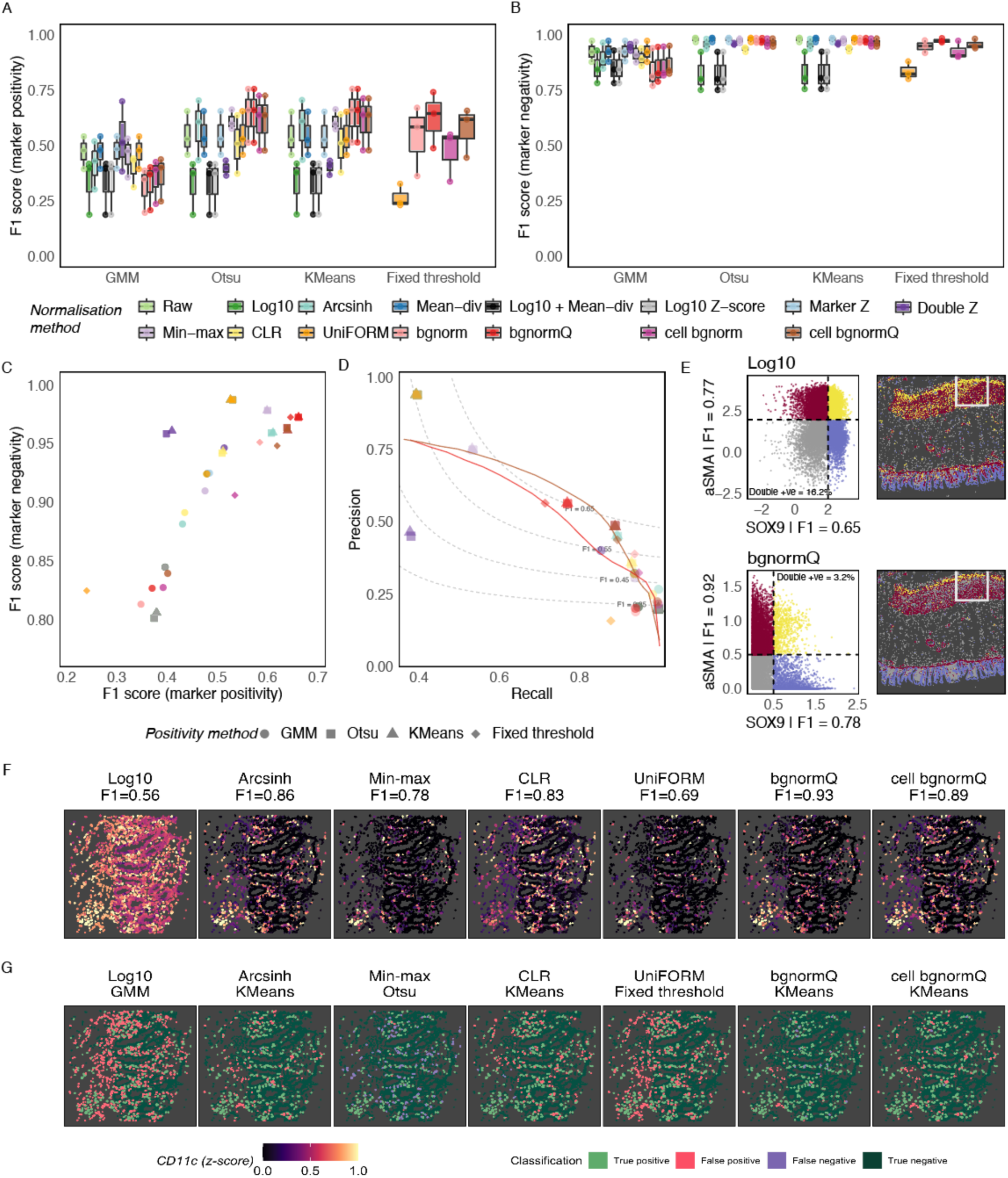
Benchmarking bgnorm for marker positivity classification shows improved performance compared to competing methods. A) Marker-specific F1 scores obtained for positivity classification using normalisation methods combined with three binary classification approaches and a fixed thresholding approach for quantile normalised bgnorm. B) Marker negativity classification F1 scores similar to A). C) Median positivity and negativity classification F1 scores per normalisation and classification method combination shows that bgnorm balances classification for both positivity and negativity while methods like UniFORM sacrifice positivity classification performance for improved negativity classification. D) Precision-recall plot with F1 score contours showing the mean precision and recall values for all pairs of normalisation and classification methods, along with the precision-recall curve for quantile-normalised variants of bgnorm as thresholds change. E) Fixed thresholding of quantile normalised bgnorm adjusted intensities retrieves the correct cell types (epithelial and stromal) with fewer double positive cells (incorrect calls), compared to a similar fixed thresholding strategy using standard log-intensities. White square marks the region with gold standard positivity annotations. F) Normalised CD11c (dendritic cell marker) obtained using different methods showing the best possible positivity classification obtained in G). F1 scores are reported for each method along with distribution of misclassified cells (false positives and false negatives).

Interestingly, a fixed threshold of 0.5 with either cell-or pixel-level quantile normalised variants of *bgnorm* (bgnormQ and cell bgnormQ in Figure 3A-E) resulted in better performance than all other normalisation methods combined with different classification approaches. The only other method designed for a fixed threshold, UniFORM, demonstrated significantly worse performance than all *bgnorm* variants. This result highlighted the importance of the model-based quantile normalisation included in *bgnorm* in enabling comparisons across samples, markers, and datasets. Intuitively, instead of moving the threshold to classify the data, *bgnorm* shifts the data to match a fixed threshold. Studying two markers of distinct cell populations, αSMA and SOX9, representing stromal and epithelial progenitor cells respectively in a healthy colon, we see how a single fixed threshold of 0.5 for quantile normalised *bgnorm* adjusted data yields better separation of cell types than a shared threshold selected to maximise separation for standard log normalised intensities (Figure 3F). Studying the precision and recall values contributing to the F1 score shows that *bgnorm* maximised precision while maintaining recall at the level of competing methods, thus offering balanced performance (Figure 3E). The only other methods with significantly higher precision (UniFORM and Min-Max) had less than half the recall rates. Since cell type identity is often determined using multiple markers, higher per-marker precision at the expense of recall is rarely preferred. However, precision still matters since finer subtyping heavily relies on the presence of a single marker. In this context, *bgnorm*’s balanced performance is beneficial for cell typing in spatial proteomic datasets. When using a fixed threshold, increasing the threshold increased the precision while sacrificing recall (Figure 3E). The threshold of 0.5 resulted in nearly optimal performance for positivity allocation for the pixel-level quantile-normalised *bgnorm* adjusted data.

While the benchmark summarises overall performance, the biological implications are not as clear. To demonstrate the impact of normalisation on biological discovery, we visualised performance on a marker showing the greatest divergence in performance across methods, CD11c. CD11c is included in most panels to study dendritic cells. Accurate identification of dendritic cells is required to study downstream immune interactions in health and disease. Visualising normalised data using different approaches showed that most methods other than log10 improve CD11c detection (Figure 3F). Assessing classifications for this marker showed that most misclassified cells following Arcsinh, CLR, UniFORM, and *cell bgnormQ* normalisation were false positives, while Min-Max normalisation resulted in a higher false negative rate. Interestingly, while false positives for other methods were diffuse in space, they were localised adjacent to true positives for *cell bgnormQ* data. This is not an issue with pixel-level *bgnormQ* suggesting that while the cell-level *bgnorm* approximation works quite well in most cases, it does not fully capture the pixel aggregation process used to generate cell-level intensities.

Like normalisation, quality control (QC) of multi-sample multiplexed imaging-based spatial proteomic datasets can be tedious. For instance, the full study of the 3D skin dataset is composed of a total of 5616 images, spanning 26 slices measured across 12 samples (labelled as Regions) and 18 proteins. Manual inspection of these images is rarely feasible, motivating the requirement of automated approaches. One aspect of quality control is the assessment of signal to noise ratio to determine whether antibody staining has worked. Since we already model intensity data by decomposing them into technical and non-technical sources of variation, we can use the existing *bgnorm* model fit to estimate the signal to noise ratio. Specifically, we can compute the divergence between the signal component (component 3) and the non-specific binding component (component 2), with increased divergence of the distributions reflecting increased signal compared to background. With this rationale, we propose a model-driven QC metric in the form of the Jensen-Shannon Divergence (JSD) measurements computed between the third and second components of the *bgnorm* model. Higher values of this metric correlate with higher signal-to-noise ratio thereby representing better quality. A value of zero represents the case where the signal is completely indistinguishable from background variation, therefore no usable information exists in the data.

Computing this QC metric for the 3D skin data shows sample, channel, and slice level variation in signal quality (Figure 4A). Defining low-quality images as those with JSD < 0.2 and moderate quality images as those with 0.2 < JSD < 0.3, we were able quickly identify QC failure in CD68 for four samples from the sample Region 8. While images stained to have moderate quality are not necessarily failure cases, these are flagged for further manual inspection. Since the proposed QC metric is dependent on the model fit, we also overlay the proportion of pixels estimated to be positive for a given protein image (π_3_). This forms a biologically informed QC metric that should be assessed in the context of the biological system being studied. For instance, the proportion of FOXP3 positive cells is unusually high in a small subset of slices from sample Region 7. FOXP3 is a marker of regulatory T cells, which are often rare under physiological conditions thus contradicting the observed proportions. Since the proportion can be higher during inflammation, these subset of images should be assessed and curated for image quality in the context of the biology.

**Figure 4:**
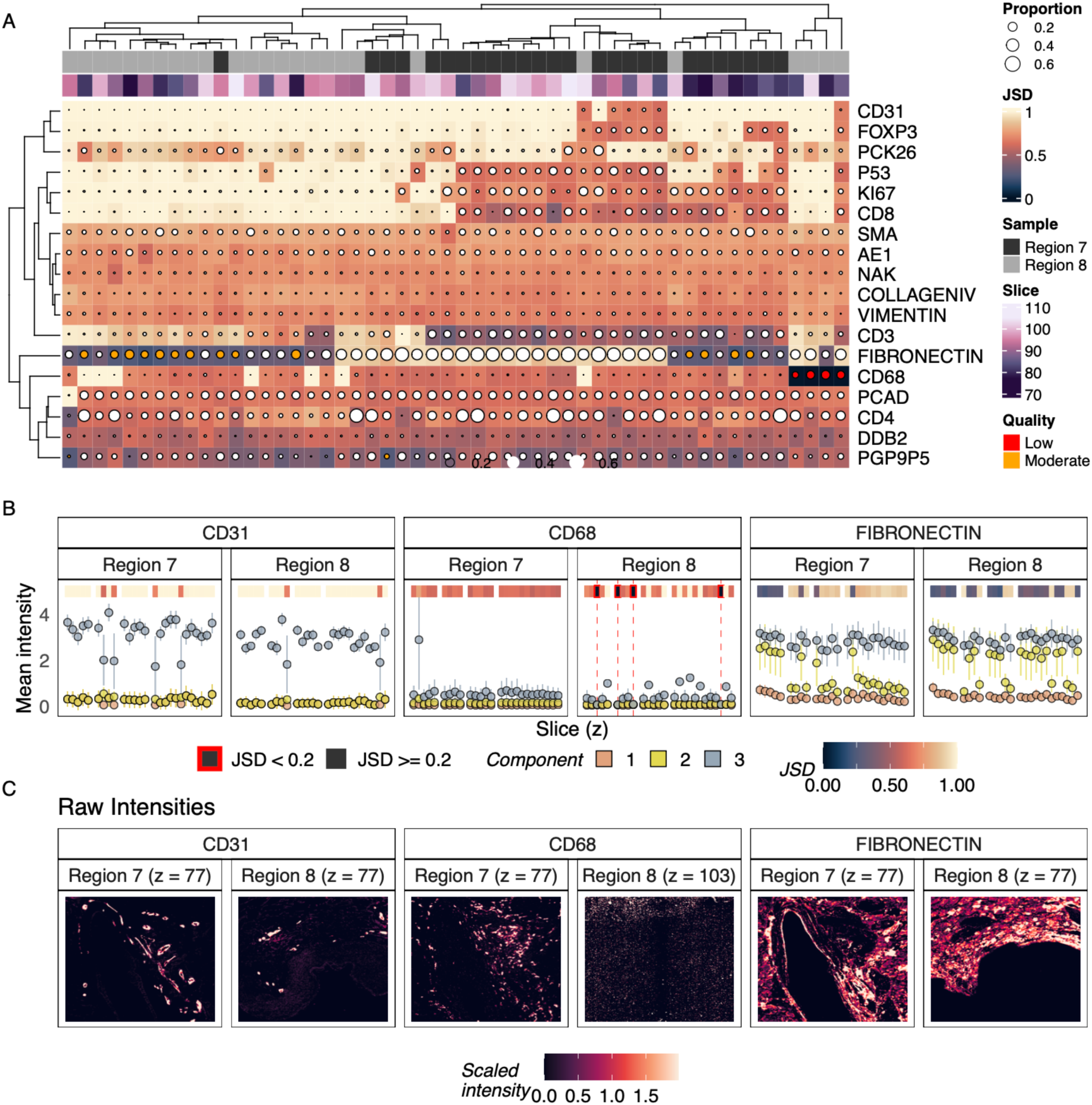
Model-based quality control of multiplexed immunofluorescence imaging data using the Jensen-Shannon Divergence (JSD) metric. A) Heatmap showing the quality metric, JSD across proteins and images, where each image represents a single z-stack slice from one of two 3D tissues studied. Circle sizes represent the proportion of pixels classified in the signal component (component 3) and are coloured red for low-quality data (JSD < 0.2) and yellow for moderate quality data (0.2 < JSD < 0.3). B) Three representative proteins selected to demonstrate high-quality measurements across all samples and slices (CD31), moderate quality but not necessarily failure cases across some slices in some images (FIBRONECTIN), and low-quality failure cases in some slices from a single sample (CD68), with JSD statistics, and component means and standard deviations visualised to demonstrate signal to background contrast. C) Representative images from each case in B) showing good (CD31), moderate (FIBRONECTIN), and poor-quality (right panel of CD68) images from each sample. For CD68, a good (left) and poor (right) quality image as determined using the JSD from each sample is shown side-by-side demonstrating that the JSD is able to identify differences in data quality.

Studying model parameters across different markers revealed the types of trends captured by the JSD. Detection quality of CD31 was among the highest, producing higher JSD values across samples and slices, and this was due to significant separation between the signal and background components (Figure 4B). Investigating the images showed a striking contrast between CD31+ and CD31-pixels (Figure 4C). A moderate JSD between 0.2 and 0.3 was observed across some FIBRONECTIN images, however, this was not due to quality issues with the images but instead a reduced signal to background ratio as assessed by the mean and variances (Figure 4B-C). Finally, the JSD for CD68 was lower than 0.2 for four slices from Region 8 indicating that the distribution of signal was not as clearly distinguishable from that of the non-specific binding. This is evidenced by the presence of bright FIBRONECTIN+ pixels with some persisting luminescence in the background tissue. A low JSD can either indicate failure in the data generation process, or the absence of signal in the tissue therefore users would have to manually assess flagged images. Overall, the QC heatmap (Figure 4A) produced using *bgnorm* can drastically reduce the degree of manual inspection required to assess quality in cohort-scale studies.

### Assessing spatial variability in intensity distributions using whole slide images

Spatial proteomic profiling is typically either performed at the whole slide level, or for tissue microarrays constructed using multiple cores. The proposed background correction approach using *bgnorm* is equally applicable to both setting with the recommendation being that individual sample be processed independently. Since tissue samples in tissue microarrays are small, spatial technical variation is expected to be minimal therefore *bgnorm* can be applied without modelling any spatial variation. The same cannot be claimed for larger whole-slide images from whole tissue profiling. The presence of spatial technical variation would result in spatially varying model components therefore resulting in biases in background correction across the tissue. To assess the presence of technical variation, we designed a simple computational experiment using a head and neck cancer dataset (Figure 5A) where we compared GMM model parameters computed across the whole tissue against tile-specific parameters computed using pixels from each tiled image (Figure 5B).

**Figure 5:**
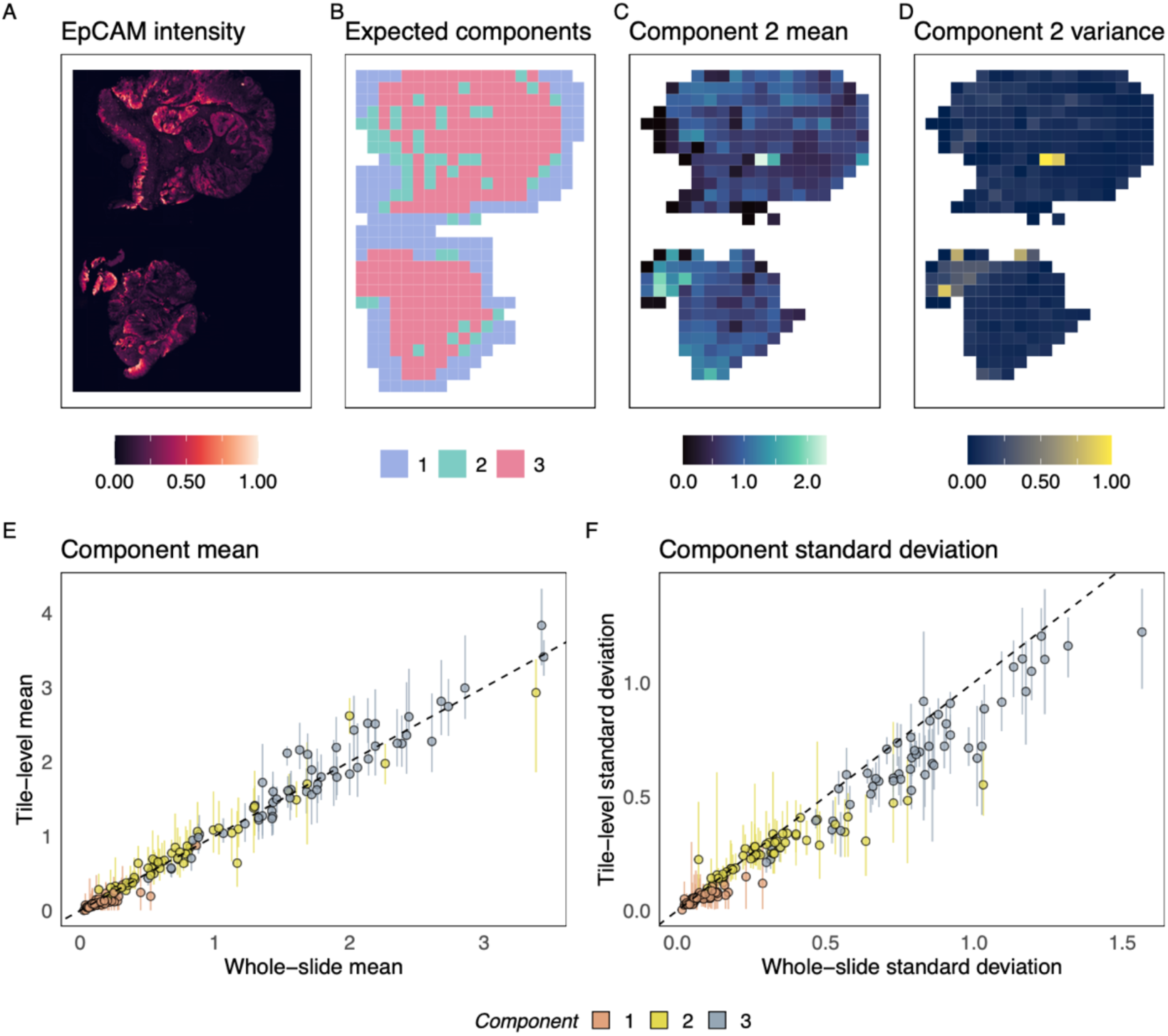
Model parameters are spatial invariable across whole-slide images negating the need for a spatially aware background adjustment model. A) Example image of a single protein, EpCAM, for a whole slide image. B) Tiling showing the expected number of components within each tile of the EpCAM image; tiles from background areas are likely to have only one component while tiles from tissue regions may have 2 or 3 components present depending on whether background tissue is captured or not. C) Component 2 means estimated across all tiles showing little spatially coherent variation in tile means. D) Variation in component 2 variances across tiles as in C). E) Comparison of tile-level and whole slide component means with the variance in tile-specific means visualised as error bars (1 standard deviation) shows equivalence in estimated parameters. F) Comparison of component standard deviations like E).

Since tissue composition varies across tiles, the true number of components for the GMM would vary across different tiles in the tissue. A three-component model would be appropriate only for tiles with the protein of interest, some background tissue, and some empty space, while a two-component model would be required if empty space was present. Where only two or one of these components are present, a two-or one-component model was used. Since this information is not known for each channel, an approximation is achieved by first fitting the three-component GMM to the full dataset, classifying pixels, and subsequently assessing the number of components present. Studying EPCAM intensity across the image showed that this strategy was able to allocate the expected number of clusters within each tile (Figure 5B). While the expected number of clusters may be incorrect for some tiles, on average the allocation works well. Studying means and variances of component 2 for the EPCAM models showed that both parameters broadly varied stochastically, with little spatially structured variation (Figures 5C-D). Some outliers were observed due to inaccurate model specification (i.e., incorrect estimation of expected components).

Finally, studying mean and standard deviation parameters for all three components across all channels showed that tile-level parameters were comparable to tissue-level estimates (Figures 5E-F). Majority of the component means and variances across channels had the tissue-level estimate within one standard deviation of the tile-specific parameters. Parameter variability was significantly lower in the background and non-specific binding components compared to the signal component, suggesting that most of the signal variation across the tissue was biologically driven. Collectively, these results suggest that: i) both spatial and non-spatial variation is minimal in the context of within-component variation across the slide, i) spatial variation is primarily biology driven where it does exist, and iii) that a spatially aware background correction model is not needed for whole-slide images. Should spatially aware modelling be required for a specific study, spatially aware normalisation models such as SpaNorm ^27^ could be applied to bgnorm adjusted data.

## Discussion

Preprocessing datasets is crucial to any analyses and can lead to downstream analytical challenges if not performed appropriately. Here we tackled the preprocessing steps of quality control and normalisation for multiplexed immunofluorescence-based spatial proteomic datasets. We proposed the three-component mixture of Gaussian convolutions generative model for imaging-based measurements and derived methods for background adjustment and quality control based on modelling parameters. While the model broadly represents the three primary sources of variation, the Gaussian assumption may fail in rare cases where the data is partly skewed, or outliers exist due to imaging artefacts or intentional masking of regions. In such cases, while not entirely accurate, a non-parametric or more powerful three component classification algorithm, such as one built using deep learning frameworks, may be used to classify the pixels, and estimated means and variances could be plugged into the deconvolution framework to obtain background corrected data. Nonetheless, where the current *bgnorm* algorithm fails to accurately estimate model parameters, downstream processing such as positivity calling can still be corrected through user intervention because *bgnorm* simply applies a continuous monotonic transformation to the data.

Experiments performed in this study primarily involved data obtained from the Akoya Phenocycler Fusion (previously CODEX) and the Cell DIVE platforms, however, the results obtained are not limited to this technology. At the core of the technology, antibodies with fluorescence tags are used to measure the abundance of proteins. In areas of the slide devoid of tissue, antibodies are unlikely to bind and there is some basal level of illumination measured due to instrument noise. Antibodies have varying binding affinities and specificities therefore we are bound to observe non-specific binding. Finally, antibodies are more likely to bind to the target protein than elsewhere. These three processes are captured by the generative model we proposed. Many technologies use a similar concept with minor variations, including but not limited cyclic immunofluorescence (CyCIF), Opal, and CosMx CellScape PSP. Additionally, while technologies like imaging mass cytometry (IMC) do not use fluorescence tags, they rely on metal-tagged antibodies which will have both target and off-target binding, as well as some basal level of measurement noise, thus making *bgnorm*-based modelling appropriate. This makes the *bgnorm* framework powerful with minor finetuning of parameters such as the cofactor required for different technologies. While foundation models may learn to account for batch effects implicitly given sufficient training data, gains performance and representation quality in such models plateau with dataset size and are not automatically robust to measurement quality^28, 29^. *Bgnorm* therefore is a potentially complementary approach, providing cleaner and higher-quality data making model pretraining and inference more efficient, with its learned representations more reflective of the underlying biology rather than technical variation.

A key strength of the *bgnorm* framework is that it does not simply rescale measurements, which is what most current normalisation methods do, but instead reduces variation from background sources of intensity. Comparability across measurements is instead performed through a subsequent quantile normalisation. As such, *bgnorm* first increases the contrast within a measurement unit (i.e., a single channel) and then enables comparability across samples, slices (in 3D images), and channels. Through an extensive benchmarking using independent gold-standard annotations, we showed that *bgnorm* without quantile normalisation performs better than all other normalisation methods that only rescaled the data. Subsequent quantile normalisation rescaled all measurements to the same scale, making the simple method of using a fixed common threshold more powerful than pairing complex classification methods with other normalisation methods. A noteworthy feature here is that though normalisation is performed within a single image, measurements are made comparable across samples, channels, and slices. This means that *bgnorm* can be applied to images sequentially in large atlas-scale studies as they are added, removing the need for renormalising the entire atlas.

Model-based quality control was proposed in this study to quantify the signal to background ratio using the Jensen-Shannon Divergence (JSD) as a metric. We showed that small values of JSD were automatically able to flag failed images, however, this needed to be coupled with biological knowledge of the system being studied to enable a more comprehensive quality control assessment. Beyond a single study, the proposed QC framework can be used to learn more about the performance of antibodies when applied across multiple datasets and tissue types. For instance, JSD metrics could be collected from all studies involving a single antibody, and the distribution could be studied to assess overall quality of the antibody across different labs, equipment, and tissue types. Specific issues could be identified by studying the individual model parameters. Alternatively, such parameters could be studied to learn about variation in different biological systems such as different tissues, organs, or cell types. Modelling-based analysis enables such studies in the long run as we learn more about the data generation process.

Overall, we provide a significant advancement in the understanding of antibody-based protein detection through the proposal and validation of a new generative model for such data. We used this modelling framework to propose normalisation and quality control methods with easy-to-use implementations in R and python. While used for preprocessing in this study, the modelling framework can be used to characterise and study many more properties of the spatial protein datasets, including technical and biological sources of variation.

## Methods

### Image Processing

Multiplex immunofluorescence (mIF) images were processed at the pixel level prior to background normalisation. Raw intensity values were first median filtered using a 3×3 median filter to reduce high-frequency noise while preserving local spatial structure within tissue regions. Intensities were subsequently log2-transformed using a fixed cofactor to reduce variation in lower intensity fluorescence signals.

The transformed intensity for a given pixel intensity *I* was defined as:

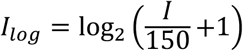

where 150 represents the cofactor used for signal compression for 16-bit images.

### The *bgnorm* model for background adjustment

Multiplex immunofluorescence (mIF) imaging data contain multiple sources of signal variation, including background noise, tissue autofluorescence, non-specific antibody binding, and true biological signal. Following log-transformation, pixel intensity distributions commonly exhibited multimodal structure, motivating the use of a probabilistic mixture modelling framework for signal decomposition. We assumed that observed pixel intensities arise from three underlying sources corresponding to:

1. Those that contain only background noise (*X*_1_ = *U*_1_)
2. Those that contain background noise and additional signal due to non-specific binding (*X*_2_ = *U*_1_ + *U*_2_)
3. Those that contain background noise, non-specific binding and true biological signal (*X*_3_ = *U*_1_ + *U*_2_ + *U*_3_)

For a particular pixel *i*, we assumed that each component (*X*_1_, *X*_2_, *X*_3_), follows a Gaussian distribution:

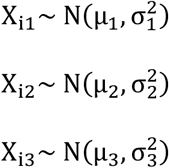

Because the true signal state of each pixel is unobserved, the marginal distribution of observed pixel intensities was modelled using a three-component Gaussian Mixture Model (GMM). The probability density function of the observed intensity distribution was therefore defined as:

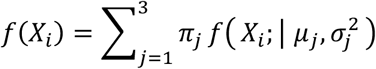

where *π*_j_, *μ*_j_, and *σ*^2^ denote the mixture proportion, mean, and variance of component *j*, respectively.

For each pixel *i*, let *C_i_* denote the Gaussian mixture component to which the pixel belongs. Here we derive a component specific background correction formula.

If a pixel belongs to the first component (*C_i_* = 1), it is assumed to contain only background signal. Consequently, the expected biological signal conditional on the observed intensity is zero:

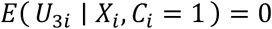

If a pixel belongs to the second component (*C_i_* = 2), it is assumed to contain background and non-specific binding signals but no true biological signal. Therefore, the expected biological signal conditional on the observed intensity is also zero:

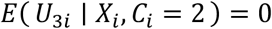

Pixels belonging to the third component (*C_i_* = 3) are assumed to contain true biological signal in addition to background-associated effects. For these pixels, we would like to retain the signal contribution arising from *U*_3*i*_ while removing the contributions arising from *U*_1*i*_ and *U*_2*i*_. This was achieved by estimating the conditional expectation of *U*_3*i*_ given the observed pixel intensity *X_i_*.

Using the standard conditional expectation formula for a bivariate normal distribution,

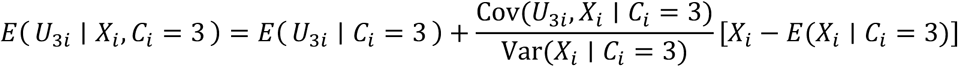

The Gaussian mixture model directly provides estimates of the conditional mean and variance of *X_i_* under the third component. To estimate the corresponding quantities for *U*_3*i*_, we note that

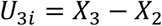

Therefore, the expected biological signal can be estimated as:

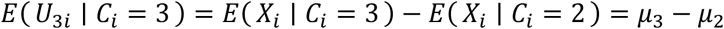

The variance of the biological signal component was estimated as:

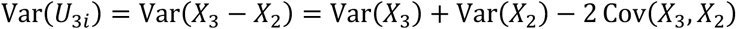

Following the original derivation, the covariance between *X*_3_ and *X*_2_ was approximated by the minimum of their variances,

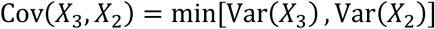

Substituting the fitted Gaussian mixture variances gives:

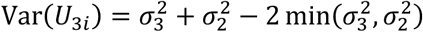

Assuming independence between *U*_3_ and the background-associated components *U*_1_ and *U*_2_,

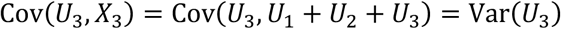

Therefore,

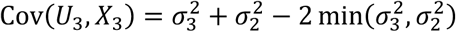

Substituting these quantities into the conditional expectation formula yields:

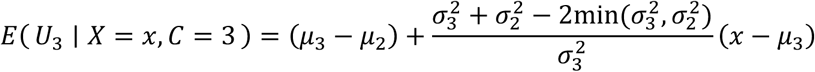

In practice, pixel identity is unknown and must be inferred probabilistically. Therefore, the final corrected intensity was calculated as the posterior-weighted average of the component-specific corrected signals:

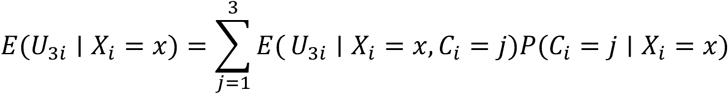

Since the expected biological signal is zero for the first two components, this expression simplifies to:

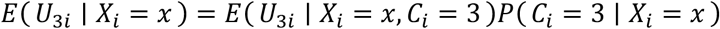

where *P*(*C_i_* = 3 ∣ *X_i_* = *x*) is the posterior probability that pixel *i* belongs to the biological signal component.

The final adjusted intensity was therefore calculated as:

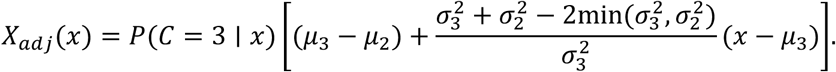

Model parameters for the GMM were estimated using the Expectation-Maximisation (EM) algorithm implemented in the *mclust* ^30^ and *sklearn.mixture* ^31^ packages for the R and python implementations respectively. When component means are close which often happens for components 2 and 3, the stochasticity induced by different initialisations of the GMM fitting procedure and data sampling can result in mislabelling of components. For instance, the mean of the true component 2 will be marginally higher than that of component 3 due to stochasticity. This is often the case when tails of component 3 are heavier than that of component 2. In such instances, we assess which component has heavier tails at quantiles larger than the 50%-tile and annotate the distribution with the heavier tail as component 3, the signal component. In all other instances, the component with the highest mean is the signal component. The deconvolution formula derived above was then applied to obtain adjusted log-intensity values.

### Cell-level background normalisation

At the cell level, aggregated intensities were assumed to primarily consist of non-specific staining/autofluorescence and true biological signal, while pure background signal was assumed to be largely absent following cell segmentation. Pixel intensities were aggregated to the cell level using the arithmetic mean of segmented pixels and subsequently log transformed using the same cofactor (150) as the pixel-level model prior to fitting the two component GMM. Consequently, cell-level intensity distributions were modelled using a two-component Gaussian Mixture Model (GMM) corresponding to:

1. Those that contain only background noise and additional signal due to non-specific binding (*X*_1_ = *U*_1_)
2. Those that contain background noise, non-specific binding and true biological signal (*X*_2_ = *U*_1_ + *U*_2_)

Model fitting was performed using the same EM based framework described for pixel-level modelling. The deconvolution-based correction framework, though being an approximation, was subsequently applied, with the second Gaussian component representing the biological signal distribution.

Following the pixel-level derivation, corrected cell-level intensities were estimated as the posterior-weighted expectation of the biological signal component:

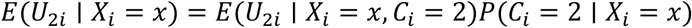

where *P*(*C_i_* = 2 ∣ *X_i_* = *x*) denotes the posterior probability that cell *i* belongs to the biological signal component.

The final adjusted cell-level intensity was calculated as:

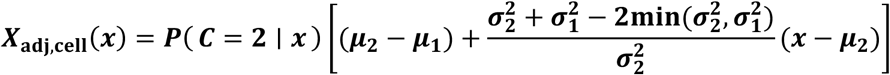

### Jensen-Shannon Divergence (JSD)

The Jensen-Shannon Divergence is a symmetric quantity to compare two distributions, derived from the Kullback-Leibler (KL) divergence. The Kullback-Leibler divergence between two normal distributions P and Q can be calculated as:

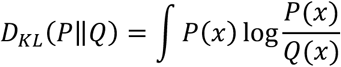

Given *M* = (*P* + *Q*)/2 is a mixture of *P* and *Q*, the JSD can then be calculated as:

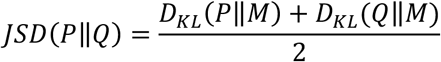

### Model-based quantile normalisation (*bgnorm^Q^*)

Conventional quantile normalisation divides each measurement by a specific quantile of the data, such as the 75%-tile. While the observed quantile suffices in most cases, we want to normalise the data by the quantile of the signal component. As such, we need to derive the normalisation factor from the quantile of the signal component. The simplest way to do so is to compute the desired quantile of the third component (e.g., *q*^0.75^, the 75%-tile of *N*(*μ_3_, σ_3_^2^*)) and then “adjusting” it using *bgnorm*. This results in the normalisation factor below:

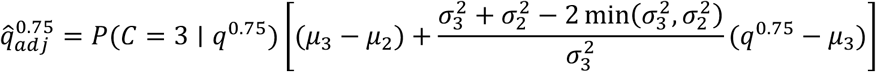

### Spatially adaptive tile-based modelling of whole-slide tissue data

Whole-slide multiplex imaging data were partitioned into a 20 × 25 grid of spatial tiles. This tiling strategy roughly reflected the stitched field-of-view (FOV) structure of the imaging acquisition, while still maintaining enough pixels within each tile for stable Gaussian mixture model fitting. A bgnorm model was fit to the whole slide image of each channel, pixels were classified into the three components, and the proportion of each component estimated within each tile. Proportions less than 0.01 were set to 0 as they could result from stochasticity. The expected number of components per tile was then the number of components with non-zero proportions (Figure 4B). A *bgnorm* model was then fit for each channel within each tile and the identity of each component was resolved on whether the original proportions were non-zero or not. For instance, if a tile had 0% pixels in the first component, 80% in the second and 20% in the third, a two component GMM would be fit to said tile and the identity of the inferred components would be components 2 and 3 representing autofluorescence/non-specific binding and signal respectively. Tile-level parameter estimates, specifically component-specific means and standard deviations, were compared with whole-slide model estimates to assess spatial stability of the inferred mixture distributions.

### Benchmarking

To assess the impact of background adjustment on downstream marker positivity classification, we benchmarked *bgnorm* against commonly used preprocessing approaches using expertly annotated Pan-M Multiplex imaging dataset ^25^. Benchmarking was performed using three CODEX (now Akoya PCF) healthy colon fields of view (FOVs) containing 10,749 unique cells and 423,036 expert reviewed marker positivity annotations. Of these, 406,352 annotations carried a strictly binary (positive/negative) gold-standard label and were used for scoring; annotations flagged as ambiguous during proofreading or lacking review were excluded. As annotations were provided at the marker level, individual cells could contribute to multiple positivity labels.

To assess the downstream impact of normalisation, we compared commonly used preprocessing approaches with our method on downstream marker positivity classification ^11^. Preprocessing methods compared included: untransformed (raw) intensities; log10 transformation; arcsinh transformation (cofactor = 150); mean normalisation (marker intensity divided by its mean); marker-wise z-score normalisation, computed both on raw intensities and on log10-transformed intensities; double z-score normalisation (sequential marker-and cell-wise z-scoring); centred log-ratio (CLR) transformation; min-max scaling (1st–99th percentile, per channel); UniFORM, a cross-sample histogram registration method¹⁷, applied here across the three FOVs treated as independent samples; pixel-level *bgnorm* and its quantile-normalised variant (*bgnorm^Q^*); and cell-level *bgnorm* and *bgnorm^Q^*, in which pixel intensities were aggregated per cell using the geometric mean (Methods: Cell-level background normalisation). Because the log transform is undefined at zero, cells with a raw intensity of exactly zero were excluded from log-based transformations (log10, log10 with mean normalisation, and log10 z-scoring) and instead called negative for that marker since the protein was undetected.

Following preprocessing using the different methods, marker positivity was assigned independently for each marker using Gaussian mixture modelling (GMM), Otsu thresholding, k-means clustering, or a fixed threshold classification. GMM was fit with two components, and the component with the higher mean was assigned as marker positive. K-means clustering was similarly performed with two clusters, and the higher centroid cluster was assigned as marker positive. For Otsu thresholding, cells with marker intensities greater than the marker specific Otsu threshold were assigned as marker positive. For pixel and cell level *bgnorm* and *bgnorm^Q^* layers, an additional fixed-threshold classifier was evaluated by sweeping thresholds from 0 to 5 in increments of 0.05; a threshold of 0.5 is reported as the fixed-threshold result given its near-optimal performance (see Results). For UniFORM, positivity was additionally assigned using the algorithm’s own global-thresholding approach: a two-component GMM was fit once to log10-transformed intensities pooled across all three FOVs after cross-FOV registration, and the resulting per-marker threshold was applied uniformly to every FOV.

Predicted positivity labels were compared with expert reviewed annotations. For each FOV, classification performance was quantified as the pooled F1 score, precision, recall, and balanced accuracy across all annotated cell-marker pairs with a strictly binary gold-standard label, computed separately for the marker-positive and marker-negative classes. Channels with no positive ground-truth cells within a FOV were excluded from positive-class pooling since the precision is always 0 and recall is undefined for these channels. All channels were retained for negative-class scoring since no channel had a 100% positivity rate. Per FOV metrics were summarised using the mean and median across the three FOVs for each combination of preprocessing method and classification approach. Precision-recall analyses were additionally performed by sweeping the fixed threshold across *bgnorm* derived layers to evaluate the trade-off between sensitivity and specificity.

## Data availability

The 3D multiplex skin imaging raw data was downloaded from the Human Biomolecular Atlas Program (HuBMAP) Data Portal (https://portal.hubmapconsortium.org/browse/collection/34b068d4a926f77fd98b3d968b6c17 <u>2f</u>) ^32^. The CODEX colon dataset used for benchmarking was obtained from the Pan-M dataset (https://huggingface.co/datasets/JLrumberger/Pan-Multiplex) ^25^. The CODEX NSCLC tissue microarray and HNSCC whole-slide tissue section datasets are available at https://zenodo.org/records/20791696.

## Ethics declarations

In this study, we identified a HNSCC patient from the Princess Alexandra Hospital (PAH) (HREC/2022/QMS/89,452), that was determined eligible for inclusion in our study. Formalin-fixed paraffin-embedded (FFPE) tissue specimens was collected. To ensure the exclusion of non-neoplastic epithelial cells, expert pathologists demarcated tumour and stromal regions. The CODEX NSCLC is comprised of a cohort that contained both squamous and adenocarcinoma NSCLC histology. This has Queensland University of Technology (QUT) Human Research Ethics Committee approval (UHREC #2000000494) and University of Queensland ratification.

## Code availability

*Bgnorm* is available via GitHub as both R (https://github.com/BhuvaLab/bgnormR) and python packages (https://github.com/BhuvaLab/bgnormPy).

## Acknowledgements

We want to thank Professor Terence (Terry) Speed, WEHI, for initial discussions on modelling intensity data, and sharing his experiences in modelling intensity data for microarrays. This work was supported by resources provided by The University of Queensland Research Computing Centre’s Bunya supercomputer ^33^, and by the supercomputing resources provided by the Phoenix High Performance Computer (HPC) service at Adelaide University.

## Funding

DDB is supported by the NHMRC EL1 fellowship GNT2034141. The study investigators DDB, CWT and AK are supported by the MRFF METASPATIAL Study (2031100). AK and RT are supported by the Queensland Spatial Biology Centre at the Wesley Research Institute, the Passe and Williams Conjoint Grant, and PA Research Foundation (PARF). AK is supported by Cure Cancer Foundation.

## Notes

https://github.com/BhuvaLab/bgnormR

https://github.com/BhuvaLab/bgnormPy

https://portal.hubmapconsortium.org/browse/collection/34b068d4a926f77fd98b3d968b6c172f

https://huggingface.co/datasets/JLrumberger/Pan-Multiplex

### Competing Interest Statement

The authors have declared no competing interest.

